## Supplementary material for "Quantifying ‘just-right’ APC inactivation for colorectal cancer initiation": Supplemetnary information

### Supplementary Figures

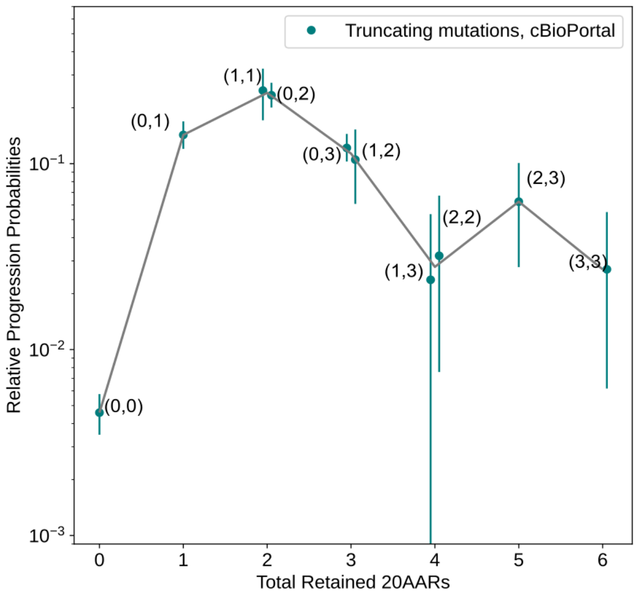

*Supplementary Figure 1. Progression probabilities of APC genotypes in cBioPortal Cohort.* The relative progression probability of different APC biallelic mutant genotypes, $\tilde{p}_{(M,N)}$, is plotted against the total number of 20AARs retained across both alleles. The frequencies of genotypes were calculated from sequence data of MSS primary CRCs in the cBioPortal cohort without copy-number alterations on APC (n=1,041, Methods). Whiskers for 95% confidence intervals (bootstrapping). The grey line is the weighted average of the progression probability over all genotypes which result in a given number of retained 20AARs.

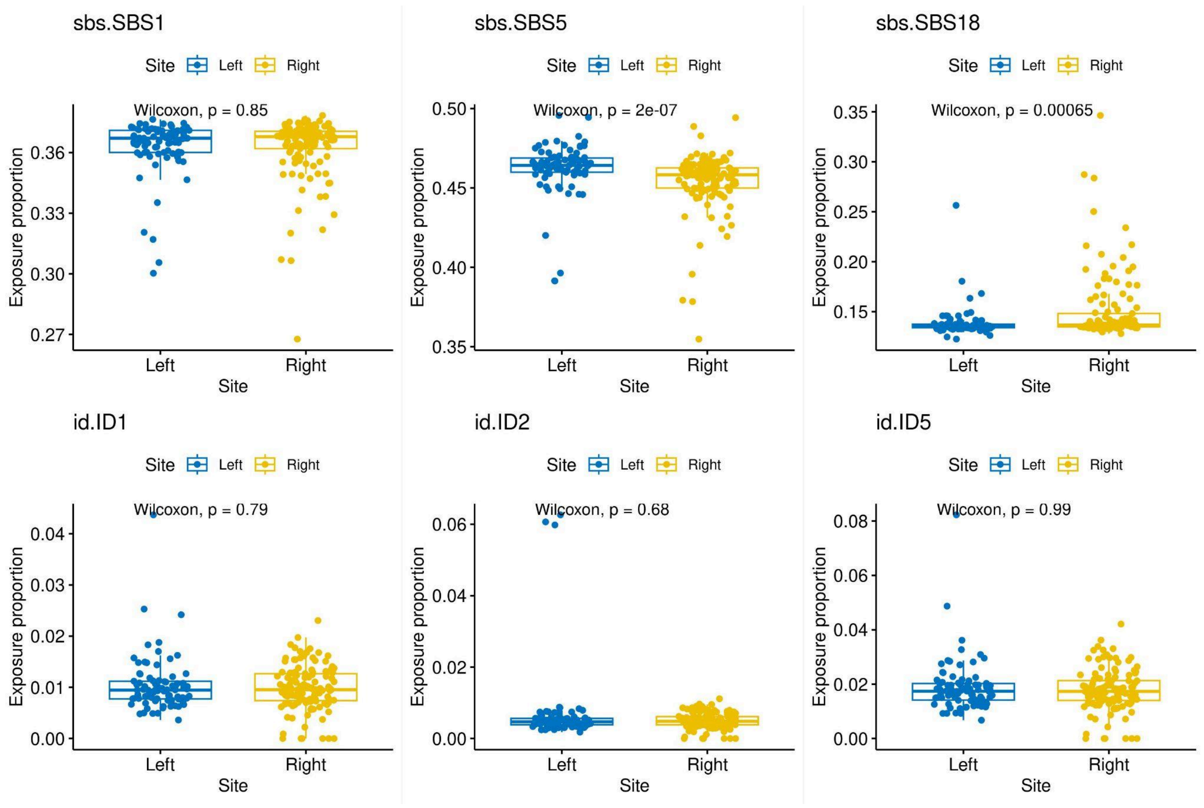

*Supplementary Figure 2. Site-specific differences in mutational signatures in the healthy colon.*

For mutational signatures observed ubiquitously in the healthy colon, we show the proportion of signature exposures, calculated by Lee-Six *et al* [2] for healthy colonic crypts labelled as Left colon (corresponding to distal) or Right colon (corresponding to proximal). Significant site-specific differences exist for SBS5 and SBS18, although the magnitude of the effect is relatively minor and so is unlikely to largely contribute to site-specificity of *APC* genotypes in CRCs.

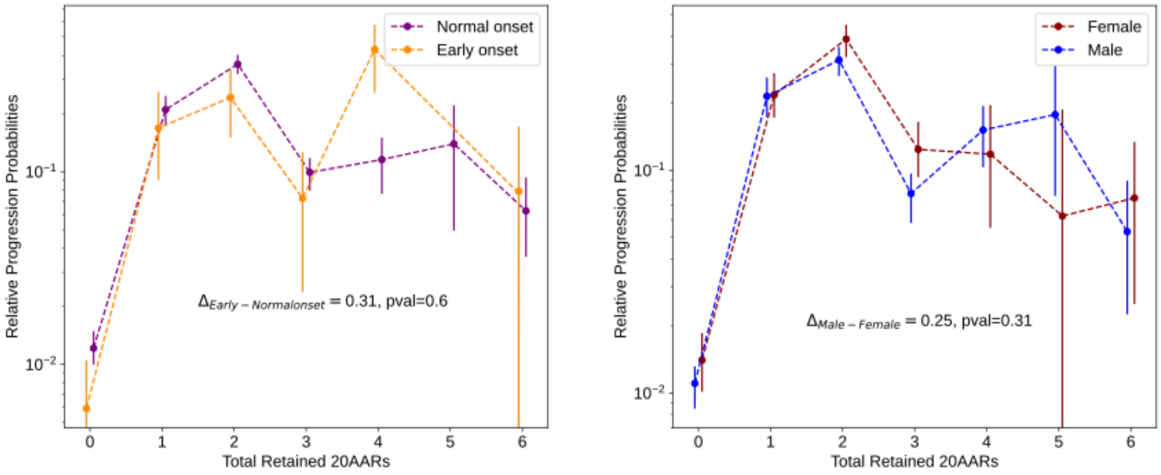

*Supplementary Figure 3. Differences in the progression probabilities by sex or age.*
The relative progression probability versus total number of 20AARs retained over both alleles, for (A) tumours in male (orange) versus female (purple) patients, and (B) in patients with early onset (<50 years old at resection, purple) versus normal onset (>50 years old at resection, yellow). Whiskers on points indicate 95% confidence intervals (bootstrapping). The difference in the progression-weighted mean 20AARs number is indicated by Δ (Methods). We find no statistically significant differences between tumours in male versus female patients (p=0.33, permutation test) nor in patients with early onset versus normal onset (p=0.73, permutation test).

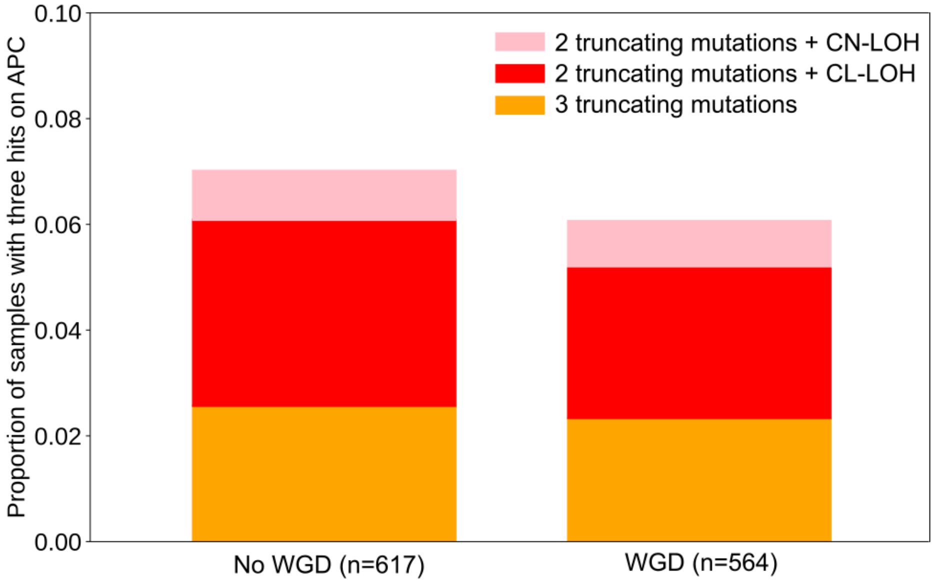

*Supplementary Figure 4. Third hits in APC in 100kGP* CRCs.

Proportion of APC-inactivated MSS primary CRCs in the 100kGP cohort (n=1,181) with evidence for third hits of different types, segregated by samples with no WGD (left) and with WGD (right), determined by Cornish et al (2022).The overall frequency of MSS CRCs with identifiable third hits in *APC* is 6.2%. Notably, in 100kGP the prevalence of third hits is not statistically different in samples with WGD p=0.58, chi2 statistic).

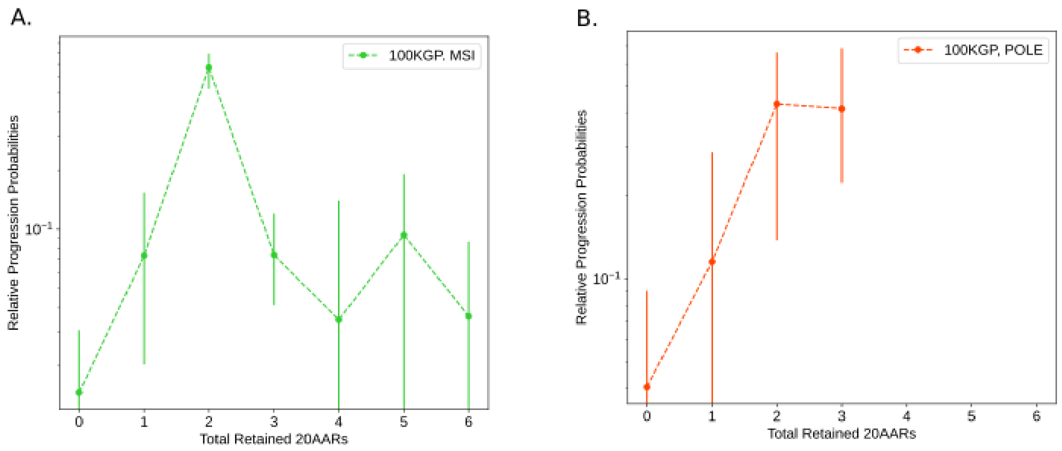

*Supplementary Figure 5. Progression probabilities in hypermutant CRCs.*
The relative progression probabilities by total number of 20AARs retained in MSI (A) and POLE-mutant CRCs (B) in the 100kGP cohort, as in main text Figure 6, but with bootstrapped confidence intervals.

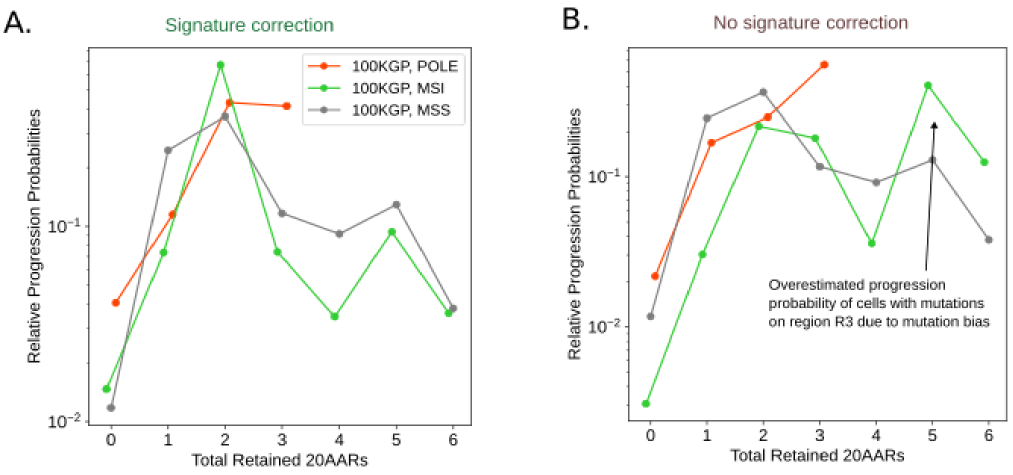

*Supplementary Figure 6. Signature correction in hypermutant CRCs.*
(A) The relative progression probabilities by total number of 20AARs retained in POLE-mutant, MSI and MSS CRCs 100kGP cohort, display overall agreement, which can be quantified by no significant differences in the progression-weighted mean number of 20AARs ($\Delta_{MSS\text{-}POLE}$=-0.22, 95% CI=[-0.62, 0.15], $\Delta_{MSS\text{-}MSI}$=-0.29, 95% CI=[-0.67, 0.08], bootstrapping). (B) As in (A) but without correcting for hypermutant mutational signatures. We seelarger differences, which can be quantified by differences in the progression-weighted mean number of 20AARs which are statistically significant in the case of MSI tumours ($\Delta_{MSS\text{-}POLE,nc}$=-0.31, 95% CI=[-0.78, 0.2], $\Delta_{MSS\text{-}MSI,nc}$=-1.87, 95% CI=[-2.51, -0.81]). We highlight that the relative progression probability is larger for 5-6 total retained 20AARs for the MSI cohort without correction, due to an indel associated mutation bias to region 3.

### Supplementary Notes

**The rate of loss of heterozygosity (LOH)**

We estimated the rates of copy-loss and copy-neutral LOH at the *APC* locus of chromosome 5 (5q22.1–q22.3) in the healthy colon by assuming phenotypic equivalence between tumours with varied molecular causes of APC loss (Methods), resulting in estimates of $5.72*10^{-6}$ and $7.18\ 10^{-6}$/cell/year, respectively.

Paterson *et al*. [3] estimated the rate of LOH through a different strategy, obtaining a higher rate of $1.36*10^{-4}$/cell/year. Paterson *et al* used the ratio of MSI cancers with two inactivating mutations in *APC* to those with one inactivating mutation and one LOH event to be 1:7, as reported by Huang *et al*. [4] based on protein assays of *n*=55 CRCs. This ratio is in high discordance with the ratio in the 100kGP cohort of 10:1, based on a large number of whole-genome sequenced samples (n=2,023) [5]. Using the inference method of Paterson *et al.* with data from 100kGP we obtained a rate of LOH of $3.89*10^{-6}$/cell/year, which is comparable to our estimate.

#### Supplementary Tables

##### Supplementary Table 1. cBioPortal datasets

| Study ID | Number of samples |
|---|---|
| coad_caseccc_2015 | 7 |
| coad_cptac_2019 | 42 |
| coad_silu_2022 | 123 |
| coadread_dfci_2016 | 105 |
| coadread_genentech | 8 |
| coadread_mskcc | 40 |
| coadread_mskresistance_2022 | 13 |
| coadread_tcga | 96 |
| crc_apc_impact_2020 | 149 |
| crc_dd_2022 | 24 |
| crc_nigerian_2020 | 11 |
| crc_public_genie_bpc | 593 |
| rectal_msk_2019 | 94 |
| Total primary CRC with two truncating mutations on APC | 1,305 |
| Total primary MSS CRC samples with two truncating mutations on APC and no copy number alterations at APC locus | 1,041 |

Supplementary Table 1. Public data accessed through cBioPortal [6,7] as of 1st of September 2023. Duplicated samples across different studies were removed. Primary tumours with two pathogenic mutations on APC and no copy-number alterations other than WGD were considered. The data and scripts used are available at https://github.com/xellbrunet/APC_Public.

**Supplementary Table 2. 100kGP APC CRC cohort**

| | Total | Primary MSS | Primary MSI | Primary POLE |
|---|---|---|---|---|
| All | 2023 | 1641 | 364 | 17 |
| APC mutant | 1499 | 1370 | 111 | 17 |
| APC biallelic inactivation | 1118 | 1037 | 64 | 17 |

Supplementary Table 2. Number of samples considered from the 100kGP CRC cohort, corresponding to that analysed by (Cornish et al. 2022).

#### Supplementary Table 3. SBS signatures in MSI CRCs in 100kGP

| Signature | Proportion of samples | Mean exposure | Mean burden |
|---|---|---|---|
| SBS1 | 0.967 | 0.127 | 12140.95 |
| SBS5 | 0.981 | 0.33 | 32135.3 |
| SBS15 | 0.368 | 0.084 | 10280.01 |
| SBS26 | 0.296 | 0.073 | 9283.43 |
| SBS44 | 0.827 | 0.31 | 34619.09 |
| SBS57 | 0.329 | 0.07 | 8026.49 |

Supplementary Table 3. Single-base substitution mutational signatures present in >20% of MSI CRCs in the 100kGP cohort, determined by [5].

#### Supplementary Table 4. ID signatures in MSI CRCs in 100kGP

| Signature | Proportion of samples | Mean exposure | Mean burden |
|---|---|---|---|
| ID1 | 0.989 | 0.134 | 18744.05 |
| ID2 | 1.000 | 0.866 | 123764.01 |

Supplementary Table 4. Insertion/deletion mutational signatures present in >20% of MSI CRCs in the 100kGP cohort, determined by [5].

#### Supplementary Table 5. Signature analysis results

| | Stop-gained | | | | | Frameshifts | | | |
|---|---|---|---|---|---|---|---|---|---|
| APC Region | Healthy colon | Healthy right colon | Healthy left colon | POLE-deficient CRCs | MSI CRCs | Healthy colon | Healthy right colon | Healthy left colon | MSI CRCs |
| *R0* | 0.848 | 0.84758 | 0.84904 | 0.78214 | 0.84827 | 0.73744 | 0.40667 | 0.50181 | 0.50368 |
| *R1* | 0.05476 | 0.05503 | 0.05438 | 0.08898 | 0.06335 | 0.06454 | 0.03586 | 0.04435 | 0.04452 |
| *R2* | 0.05457 | 0.05465 | 0.05444 | 0.04064 | 0.05034 | 0.08539 | 0.03714 | 0.04542 | 0.04558 |
| *R3* | 0.04250 | 0.04274 | 0.04215 | 0.08825 | 0.03804 | 0.11263 | 0.52032 | 0.40843 | 0.40622 |

Supplementary Table 5. The relative proportion of stop-gained and frameshifts expected to fall in different regions of *APC*, estimated by considering the ubiquitous mutational signatures found in healthy colon crypts [2], crypts with POLE mutations [8] and MSI CRCs in the 100kGP cohort (Methods). THese are used to estimate the mutation probabilities of different APC genotypes.

#### Supplementary Table 6. APC genotype mapping

| Protein position of clonal frameshift or stop-gained mutations upstream codon 1569, with i<j<k | | Copy number at APC locus | WGD | Inferred mutant type at initiation | Inferred APC genotype at initiation<br>M: region of position i<br>N: region of position j | n | Comments |
|---|---|---|---|---|---|---|---|
| i,j | | [1,1] | False | Bi-allelic mutant | (M,N) | 232 | |
| i,j | Variants need to be different | [a,b] , b>0 | True | Bi-allelic mutant | (M,N) | 187 | |
| i | | [2,0] | False | Copy-neutral LOH | (M, x2) | 104 | |
| i | | [1,0] | False | Copy-loss LOH | (M, 0) | 127 | |
| i | | [a,0], a>2 | True | Copy-neutral LOH | (M, x2) | 102 | |
| j | | [1,0], [2,0] | True | Copy-loss LOH | (M, -) | 150 | |
| i,j,k | Consider only the two most upstream mutations | [1,1] | False | Bi-allelic mutant | (M,N) | 16 | Three hits |
| i,j | Consider only the most upstream mutation | [2,0] | False | Copy-neutral LOH | (M, x2) | 1 | Three hits |
| i,j | Consider only the most upstream mutation | [1,0] | False | Copy-loss LOH | (M, -) | 5 | Three hits |
| i,j,k | Consider only the two most upstream mutations | [1,1] | True | Bi-allelic mutant | (M,N) | 13 | Three hits |

| | | | | | | | |
|---|---|---|---|---|---|---|---|
| i,j | Consider only the most upstrea m mutation | [2,0] | True | Copy-neutral LOH | (M, x2) | 5 | Three hits |
| i,j | Consider only the most upstrea m mutation | [1,0] | True | Copy-loss LOH | (M, -) | 16 | Three hits |
| i | | [a,b] with b>0 | Either | Single mutant | Excluded | 138 | |
| - | | Copy-number unknown or [0,0] | | | Excluded | 6 | |

Supplementary Table 6. Classification of *APC* genotypes at initiation. Mapping from *APC* sequence data acquired at tumour sample to *APC* genotype at initiation for all considered combinations of copy number, WGD and annotated variants.

Examples:

| Protein position of unique clonal frameshift or stop-gained mutations upstream codon 1569 | Copy number at APC locus | WGD | Inferred mutant type at initiation | Inferred APC genotype at initiation | Comments |
|---|---|---|---|---|---|
| [234 frameshift, 1311 stop-gain] | [1,1] | True | Bi-allelic mutant | (0,1) | |
| [1425 stop-gain, 1425 stop-gain] | [1,1] | True | Excluded | - | No evidence of biallelic loss |
| [1425 stop-gain] | [2,0] | Flase | Copy-neutral LOH | (2,x2) | |
| [1425 stop-gain] | [2,0] | True | Copy-loss LOH | (2,0) | |

#### Supplementary Table 7. Model parameters

| Genotype $(M, N)$ | Total retained 20AARs | Mutation probability $m_{(M,N)}$ | Frequency in 100kGP $f_{(M,N)}$ | Relative CRC progression probability $\tilde{p}_{(M,N)}$ | 95 % CI for $\tilde{p}_{(M,N)}$ | |
|---|---|---|---|---|---|---|
| (0, 0) | 0 | 0.1733 | 0.0318 | 0.0036 | 0.0024 | 0.0050 |
| (1, 1) | 2 | 0.0009 | 0.0135 | 0.1899 | 0.0962 | 0.2868 |
| (2, 2) | 4 | 0.0012 | 0.0039 | 0.0425 | 0.0000 | 0.0941 |
| (3, 3) | 6 | 0.0014 | 0.0029 | 0.0373 | 0.0000 | 0.0841 |
| (0, 1) | 1 | 0.0255 | 0.1148 | 0.0870 | 0.0687 | 0.1070 |
| (1, 2) | 3 | 0.0022 | 0.0048 | 0.0573 | 0.0170 | 0.1011 |
| (1, 3) | 4 | 0.0023 | 0.0010 | 0.0158 | 0.0000 | 0.0406 |
| (0, 2) | 2 | 0.0293 | 0.1794 | 0.1164 | 0.0943 | 0.1386 |
| (2, 3) | 5 | 0.0027 | 0.0048 | 0.0411 | 0.0136 | 0.0755 |
| (0, 3) | 3 | 0.0314 | 0.0781 | 0.0531 | 0.0407 | 0.0660 |
| (0,-) | 0 | 0.3318 | 0.0723 | 0.0037 | 0.0027 | 0.0048 |
| (1,-) | 1 | 0.0244 | 0.0897 | 0.0654 | 0.0495 | 0.0833 |
| (2,-) | 2 | 0.0281 | 0.1389 | 0.1010 | 0.0817 | 0.1231 |
| (3,-) | 3 | 0.0301 | 0.0289 | 0.0168 | 0.0107 | 0.0236 |
| (0, x2) | 0 | 0.2525 | 0.0530 | 0.0037 | 0.0026 | 0.0048 |
| (1, x2) | 2 | 0.0186 | 0.1331 | 0.1263 | 0.1016 | 0.1548 |
| (2, x2) | 4 | 0.0214 | 0.0366 | 0.0289 | 0.0185 | 0.0399 |
| (3, x2) | 6 | 0.0229 | 0.0125 | 0.0104 | 0.0049 | 0.0163 |

Supplementary Table 7. Model parameters for APC-driven CRC initiation in the healthy colon. Mutation probabilities $m_{(M,N)}$ calculated using the mutational signatures ubiquitous to colonic crypts [2], Methods), frequencies of CRCs $f_{(M,N)}$ are calculated from the 100kGP cohort of MSS primary CRCs [5], relative progression probabilities $\tilde{p}_{(M,N)}$ are calculated using $m_{(M,N)}$ and $f_{(M,N)}$ as outlined in Methods. 95% CI obtained by bootstrapping (1,000 iterations).

#### Supplementary Table 8. Odds ratio between APC and Wnt regulators

| Secondary Wnt | Odds ratio | p-value | Number of samples | | | |
|---|---|---|---|---|---|---|
| | | | APC only | Secondary Wnt only | Both | Neither |
| RNF43 | 0.019 | 3.938 E-24 | 1375 | 34 | 4 | 226 |
| ZNRF3 | 0.124 | 0.03 | 1377 | 3 | 2 | 257 |
| CTNNB1 | 0.152 | 1.464 E-05 | 1368 | 13 | 11 | 247 |
| AXIN1 | 0.281 | 0.180 | 1376 | 2 | 3 | 258 |
| AXIN2 | 0.788 | 0.590 | 1358 | 5 | 21 | 255 |
| BCL9L | 0.922 | 0.862 | 1325 | 11 | 54 | 249 |
| JUN | 1.037 | 1 | 1368 | 2 | 11 | 258 |
| FBXW7 | 1.441 | 0.155 | 1224 | 21 | 155 | 239 |
| BCL9 | 1.329 | 0.688 | 1337 | 6 | 42 | 254 |
| TCF7L2 | 2.355 | 0.001 | 1216 | 14 | 163 | 246 |
| SOX9 | 2.617 | 0.000720 | 1224 | 12 | 155 | 248 |
| AMER1 | 15.528 | 2.344 E-05 | 1301 | 1 | 78 | 259 |

Supplementary Table 8. Odds ratio and number of samples between the number of MSS primary CRCs with APC mutations and mutations in secondary Wnt drivers. P-values determined using Fisher's test. Presence of clonal driver mutations on other Wnt genes (AMER1, AXIN1, AXIN2, BCL9, BCL9L, CTNNB1, FBXW7, JUN, RNF43, SOX9, TCF7L2, ZNRF) as previously determined by Cornish *et al.* [5].

**Supplementary Table 9. Secondary Wnt variants**

| Wnt regulator | Frameshift | Stopgain | Nonsynonymous SNV | Non frameshift insertion/deletion |
|---|---|---|---|---|
| AMER1 | 4 | 38 | 3 | |
| TCF7L2 | 44 | 13 | 35 | 2 |
| SOX9 | 50 | 21 | 10 | 3 |
| BCL9 | 13 | | 6 | 14 |
| FBXW7 | 5 | 16 | 72 | 1 |
| BCL9L | 10 | 9 | 11 | |
| JUN | 4 | 2 | 2 | |
| AXIN1 | 1 | | 7 | |
| AXIN2 | 9 | 3 | 10 | 1 |

Supplementary Table 9. Number and types of variants included in the analysis of secondary Wnt regulators as previously determined by Cornish *et al.* [5].
